# Systematic standardization of organ-on-a-chip under controlled flow conditions: use case with Caki-1 and A549 cell lines

**DOI:** 10.1101/2024.10.26.619288

**Authors:** Andrea Balsa-Díaz, Laura Vázquez-Vázquez, Elías Ferreiro-Vila, Lois Rivas-Meizoso, Antón L. Martínez, José Manuel Brea, María Isabel Loza, Santiago Pérez Rodriguez, Esteban Sinde, Ezequiel Alvarez, Bruno K. Rodiño-Janeiro

## Abstract

New *in vitro* models are an urgent need for to improve both the research data and the preclinical development of new drugs. Current standardized cellular models are mainly based in 2D cell culture, which lacks flow conditions and with complex co-culture settings. In this way, advanced cell culture models, such as organ-on-a-chip (OoC), aim to solve these limitations. OoC systems are composed by a microfluidic chip functionalized with different combinations of extracellular matrixes, coatings and cell cultures to mimic the physiological conditions of human organs. Advantages of OoC include the possibility to add 3D structures, delimited regions for co-culture and dynamic flow conditions to cell cultures. However, to perform reproducible and controlled experiments with OoC, it is necessary to systematically standardize the cell culture conditions in the microfluidic channels. For this is necessary to test both the combination of flow and extracellular matrix (ECM) coating to reliably mimic the human organ physiology. In this work, we standardized both conditions, ECM coating and the flow conditions to functionalize OoC with cell lines from kidney (Caki-1) and from lung (A549) to develop OoC systems beyond the Vessel-on-a-chip setting. In this way, the protocol detailed in this work will allow to standardize cell culture on different optimized OoC types with different cell types from different origins.

## Introduction

New *in vitro* models are an urgent need to improve both the research data and the preclinical development of new drugs. Models in multiwell 2D plates have been very useful for almost 100 years (1), but this cell culture conditions are not optimal to obtain reliable biological data. Recently, alternatives for these traditional cultures have been developed, including advanced 3D cellular models classified in organoids, spheroids and organ-on-a-chip (OoC). Among these models, OoC technology has emerged as a revolutionary approach in biomedical research, being sophisticated 3D models that mimics the physiological tissular structure, enabling the generation of customized cellular microenvironments with precise fluidic and structural control (2–4).

These models emerge as an improvement of both conventional 2D cell culture (5) and animal models that, while widely used, often they generate results that cannot be directly translated to humans (6–8). Respect to the 2D cell models, OoC differ in biological response such as low cellular migration and division (9,10), cellular cycle and metabolism regulation pathways (10) and, relevantly, better mimicking the response to treatments (11,12). More specifically, Lanz *et al*. observed differential response of both 2D and 3D models are to drugs in triple negative breast cancer (11), pointing out the importance of 3D cultures as models closer to *in vivo* results and with more accurate selection of drug for each patient. Importantly, OoC can solve the current state of experimental animal use, addressing the inter-species differences and ethical issues (13). The OoC have clear advantages in the high control over the cellular environment, reproducibility and easy assembly for ease of use. The combination of the advantages of *in vitro* models and the physiological complexity (14) has risen the interest to apply OoC as preclinical in vitro models, being systems with high control over the cellular environment, reproducibility and easy assembly for ease of use. This new technology will change the process of drugs testing, allowing to improve the toxicity and efficacy testing of drugs and in the study of the pathophysiology of organs and tissues (6). In this sense, several OoC disease models have been developed, which support the idea of OoC applied to preclinical drug development phases beyond animal models for human diseases (15).

The flow within the chip is a relevant factor since it reproduces wall shear stress conditions, which are translated into biological response that affect cell differentiation, morphology and metabolism (16), being a closer mimic for real physiological environment (17). The type and the pattern of the flow are dependent of both the chip geometry and the source of external perfusion, which range from pumps, hydrostatic pressure and laboratory shakers, creating different types of flow, and subsequently, different cells’ responses (18). A traditional example of the relevance of flow over cellular phenotypes is the case of endothelial cells, where shear stress cause significant changes in cell morphology, inflammatory profile and their biomechanical responses (21), leading to the susceptibility to vascular diseases. This fact is reflected in the relation of disturbed flow (low local flow velocity zones and wall shear stress) (19) in the outer walls of bifurcation angles lead to an altered endothelial gene expression, cytoskeletal arrangement, leukocyte adhesion, oxidative stress and inflammation observed in atherosclerosis (20). However, the key role of cell culture under flow is not limited to the endothelium, since other organs, such as bone or intestine, are also conditioned by wall shear stress. Specifically, flow induced changes in osteoblast cytoskeleton and cell morphology, but also in nucleus volume and gene expression by epigenetic regulation, showing that flow can induce changes in bone development (19). In the same way, it leads to changes in Caco-2 cells to where the cell polarization is faster under microfluidic conditions than with static cultures (16).

Other relevant factor to mimic the microenvironment and to allow cells to withstand flow are the characteristics of the surface of adhesion. Polydimethylsiloxane (PDMS) is a wide use material to build OoC due to several attractive characteristics, such as its biocompatibility, gas permeability and good mechanical and optical properties (21). However, this polymer is inherently hydrophobic, leading to a poor adhesion capacity. In this step, biological functionalization has a central relevance, being extracellular matrix (ECM) coatings the most used strategy to improve the surface hydrophilicity (22) and biological motifs. Different ECM coatings can affect the cellular response (23), leading to differences in cell adhesion, biomarker production, growth rate, morphology and tight junction protein expression (24,25).

Since OoC offer promising applications in drug research for testing the efficacy and toxicity of compounds, it is essential to develop reliable and accurate *in vitro* models. The characterization of both the flow conditions and the optimized coating in new OoC will allow to obtain high quality and reproducible models for human organs and diseases. In this work, we carried out the experimental standardization of flow parameters and ECM coating using two cell lines from lung (A549) and kidney (Caki-1), being able to establish optimized conditions to ensure the reliability and consistency for OoC models.

## MATERIALS

### REAGENTS

- A549 cell line (Elabscience, cat. no. CL-0016)
- Caki-1 cell line (Elabscience, cat. no. CL-0052)
- Collagen type I (indicated as collagen 1), rat tail 3 mg/mL (Gibco, cat. no. A1048301)
- Collagen type I (indicated as collagen 2), rat tail 3-4 mg/mL (Corning, cat. no. 354236)
- Gelatine from bovine skin (Sigma-Aldrich, cat. no. G9382)
- Fetal bovine serum (FBS), Qualified (Gibco, cat. no. 10270-106)
- Fibronectin bovine plasma 1 mg (Sigma-Aldrich, cat. no. F1141)
- Ham’s F12 culture medium w/L-Glutamine (Biowest, cat. no. L0135-500)
- Hoechst 93342 (NucBlue™ Live Cell Stain ReadyProbes™ Reagent, Invitrogen, cat. no. R37605)
- McCoy’s 5A culture medium w/L-Glutamine (VWR, cat. no. 392-0420)
- Paraformaldehyde, 4% in PBS, ready-to-use fixative (Biotium, cat. no. 22023). *!CAUTION: Paraformaldehyde is toxic. Use in fume hood and wear personal protection, such as gloves and goggles*.
- Penicillin-streptomycin solution 6,0/10,0 g/L 100X (P/S, Biowest, cat. no. L0022-100)
- Poly-L-lysin solution 0.1% (w/v) in H2O (ChemCruz, cat. no. sc-286689)
- Saline phosphate buffer (PBS) tablets (Invitrogen, cat. no. 003002)
- Trypsin-EDTA 0.25% (w/v) (1X) (Gibco, cat. no. 25200-056)

### EQUIPMENT

- 3-stop platinum-cured silicone tubing 1mm ID 3mm OD (Darwin Microfluidics, SE-TUB-SIL-SSS-1*1)
- Cell culture maintenance incubator (37 °C, 5% CO2, humidified; Eppendorf CellXpert)
- Cell culture test incubator (37 °C, 5% CO2, humidified; Sanyo MCO-19AIC(UV))
- Inverted microscope (Olympus IX73P2F)
- Male luer lock connector (BFlow, B02_0027)
- Multi-linear channel chip - Set of 3 (BFlow, B06_0024)
- Open jaw slide clamp (BFlow, B02_0018)
- Reglo independent channel control (ICC) peristaltic pump (Ismatec, cat. no. MFLX78018-24-EU)
- Reservoirs with septum cap (ThermoScientific, cat. no. 047422-4001)
- Straight chip connector (BFlow, B02_0021)
- Straight press-in plug (Set of 10, BFlow, B02_0029)
- Syringe needle (BFlow, B02_0024)
- Silicone tubing 1mm ID 3mm OD (BFlow, B02_0009)
- Tubing connector (BFlow, B02_0005)

### REAGENT SETUP

#### Gelatine solution (0.02% and 0.2% w/v)

weight out the appropriate amount of gelatine for each percentage solution and dissolve in distilled water (for example 20 mg in 100 mL of distilled water for 0.02% w/v). The solution is autoclaved at 121°C for 20 minutes, allowing the complete solubilization.

#### Gelatine solution (0.02% and 0.2% w/v) with fibronectin (FN) 5 μg/mL

add the appropriate amount of fibronectin to obtain 5 μg/mL in gelatine solution 0.02% in distilled water and proceed the same way with gelatine solution 0.2%.

#### Collagen 100 μg/mL in water or PBS

add the appropriate amount of collagen to obtain 100 μg/mL in distilled water (collagen in water) or 1X PBS (collagen in PBS).

#### Collagen 100 μg/mL in water with fibronectin (FN) 5 μg/mL

add the appropriate amount of fibronectin to achieve 5 µg/mL in collagen 100 µg/mL in distilled water.

#### Staining solution

prepare a solution in serum-free medium with NucBlue 2 drops/mL.

#### Poly-L-lysin Solution 0.1% (w/v) in water

the solution is not sterile and it must be filtered before use.

#### Saline phosphate buffer (PBS)

dissolve each tablet of PBS 100 mL of distilled water. PBS solution was autoclaved at 121°C for 20 minutes for sterilization.

*Note: collagen solutions prepared in this protocol were used for channel functionalization, not for hydrogel preparation*.

### PROCEDURE

In this example, multi-linear channel chips have been used, but the same approach can be used in other OoC systems.

## 1. CELL MAINTENANCE

A549 and Caki-1 cell lines were used to perform both matrix and flow standardization. All culture media were supplemented with fetal bovine serum (FBS, 10%) and penicillin-streptomycin (P/S, 1%). Trypsinization has been inactivated using complete medium in all cell lines. A549 cells were passaged at confluence using 0.25% (w/v) trypsin-EDTA for 2 min, recovered by centrifugation at 300g for 3 min, split at a 1 plate to 4 ratio and maintained in culture in Ham’s F12 medium w/L-Glutamine (146 mg/L) with 10% (v/v) FBS and 1% P/S. Caki-1 cells were passaged at confluence using 0.25% trypsin-EDTA for 2 min, recovered by centrifuging at 300g for 3 min, split in a 1 plate to 6 ratio and maintained in culture in McCoy’s 5A w/L-Glutamine (146 mg/L) with 10% FBS and 1% P/S.

## 2. MATRIX STANDARDISATION

### 2.1 FUNCTIONALIZATION PROCEDURE. *TIMING 30 min*

2.1.1 Prepare the coating solutions to be tested (described at REAGENT SETUP section).
2.1.2 Place the autoclaved chip in a petri dish to improve handling.
2.1.3 Wash the channels with 1X PBS twice, before functionalization. *SEE TROUBLESHOOTING*
2.1.4 In each channel, place the desired coating solution by pipetting. Each channel of multi-linear channel chips (BFlow, B06_0024) holds approximately 80 µL.
2.1.5 Cover the inlet and outlets with straight press-in plug and incubate in the cell culture maintenance incubator for 16-18 hours.
2.1.6 *PAUSE POINT*: The functionalized chip can be stored in the fridge for a couple of days. It is important to seal the petri dish with parafilm to prevent dehydration. Additionally, you can place small reservoirs with water or PBS on the plate further prevent dehydration.

### 2.2 CELL SEEDING *TIMING 5h*

2.2.1 Confluent cells are trypsinised and counted using a Neubauer chamber.
2.2.2 The required cell concentration for chip seeding is 1.5·10^6^ cells per millilitre.
2.2.3 Remove the chip from the maintenance incubator and take out the straight press-in plug from the channels. *SEE TROUBLESHOOTING*
2.2.4 Remove the coating solutions from the channels.
2.2.5 Fill the channels with the cell suspension from step 2.2.2.
2.2.6 Place straight press-in plugs in the inlet and in each outlet. Remove the overflowed medium from the channels when plugs are inserted.
2.2.7 Incubate upside down for 4 hours to allow the cell to cover the top part of the channel.
2.2.8 After 4 hours, take out straight press-in plugs and repeat the seeding process (from 2.2.5 to 2.2.6).
2.2.9 Incubate the channels face up for 16-18 hours, to allow the cells to cover the bottom part of the channel.
2.2.10 Incubate the cells as long as required according to the needs of the experiment, the medium should be renewed daily.

### 2.3 LIVE CELL STAINING *TIMING 30 min*

2.3.1 After the incubation of cells for the selected experimental time, remove the chip from the maintenance incubator and take out the straight press-in plug from the channels.
2.3.2 Add the labelling solution to each channel and incubate for 30 minutes at room temperature and in the dark.
2.3.3 After 30 minutes remove the staining solution, replace with normal medium and place straight press-in plugs in the inlet and in each outlet.

### 2.4 IMAGE ACQUISITION

Measurements of the number of cells after staining were performed on an Operetta CLS High Content Analysis System (Revvity, serial number 1600L21508). Both image acquisition and analysis were fully automated using Harmony v4.9.2137.273 software. For image analysis an algorithm has been constructed in which raw images were collected, without brightfield correction, and integrate the fifty planes of each field in the same image. Next, the nuclei are specified using the B protocol predetermined by the program, which is the most suitable for interpreting images with quite separate nuclei but which may have some background. Then, the area and roundness of the cells are determined, and artifacts are excluded. Finally, the program is specified to provide the number of real nuclei (excluding artifacts). Fourteen fields are selected to cover the complete channels of the chip.

For A549 cells, 50 planes separated by 5 µm starting at 500 µm in height (up to 795) have been recorded. In the case of the Caki-1 cell line, 50 planes separated by 5 µm starting at 600 µm in height (up to 895) were also recorded. A total of 14 fields were captured using a 5x objective, obtaining images of 1080px X 1080px. The resolution is 2.39 µm per pixel. That is, the images are squares of 2581 µm on each side. All the fields from the same experiment were merged using ImageJ (v1.54f). Images were obtained using the following fluorescence channels:

| Channel | Ex/Em (nm) | Exposure time (ms) | Intensity (from maximum) |
| --- | --- | --- | --- |
| Hoechst 3342 channel (NucBlue) | 355-385/430-500 | 10 | 20% |
| Brightfield | -/655-760 | 1 | 20% |

#### BOX 1 | Fixation

1. After the image acquisition step the cells can be fixated inside the channel.
2. Take out the Straight Press-in plug from the channels and remove the medium.
3. Wash the channels with 1X PBS twice.
4. Add paraformaldehyde fixation solution and incubate for 20 minutes. *!CAUTION: Paraformaldehyde is toxic. Use in fume hood and wear personal protection, such as gloves and goggles*.
5. After 20 minutes remove the fixation solution and wash the channels with 1X PBS twice.
6. Fill the channels with 1X PBS and cover them with Straight Press-in plug.
7. Now you can store your chips until the analysis. The chips must be protected from light to avoid fluorescence bleaching.

*Note: it is important to note that viability stains will be lost with fixation*.

## 3. FLOW STANDARDIZATION

### 3.1 FUNCTIONALIZATION OF THE CHANNELS *TIMING 15 mins*

3.1.1 Coating matrixes selected from the best results from **1. Matrix standardization** were applied in the flow standardization.
3.1.2 Prepare the desired coating matrixes (described in the reagent setup). For the functionalization of a single chip, 500 µL of the matrix solution must be enough.
3.1.3 For the functionalization of the channels, follow steps from 2.1.2 to 2.1.6.

### 3.2 CELL SEEDING *TIMING 5h*

3.2.1 To culture the cells, follow the steps from 2.2.1 to 2.2.8.

### 3.3 FLOW SET UP *TIMING 1h*

3.3.1 All the elements used for the assembly of the flow (Figure 2A, Figure 2B) must have been previously autoclaved and all montage must be performed in a laminar flow hood under sterile conditions.
3.3.2 For each flow channel, two tubes must be fitted with a male luer lock connector and a syringe needle at one end and a straight chip connector at the other end (Figure 2B). One of those tubes must be a tube with stoppers to be placed in the peristaltic pump (Figure 2B). For convenience, this step can be carried out before sterilisation.
3.3.3 Place medium, previously warmed, in each of the reservoirs. You must use one vial for each flow channel.
3.3.4 Place the syringe needle end of both tubes through the septum cap of the reservoir, one inside of the medium and the other not reaching the medium (placed in the air chamber of the half full reservoir). This set-up is specified in Box 2 and it will work as a bubble trap. *!CAUTION The straight chip connector and the ends of the tubes must be kept sterile*.
3.3.5 Place the tubing with the stoppers in the peristaltic pump cassette.
3.3.6 The tube that goes from the vial to the chip must be primed. Activate the pump and let the medium flow through, when the medium starts to come out of the end of the tube, stop the pump. Then place an Open Jaw Slide Clamp near the connector end of the tube. Repeat this step with all the tubes to be used into the inlet.
3.3.7 When all the tubes are in place and primed, the chip can be connected. Remove the plugs from the first channel and connect the straight chip connector end of the tube, which comes from the vial, to the first channel inlet. Then, connect the other straight chip connector end of the tube, which goes to the pump, to the outlet (Figure 2C). The process is repeated for each channel in the chip.
3.3.8 Once all the channels are connected to the tubes, remove the open jaw slide clamps.
3.3.9 Turn on the pump and check that the flow is properly established, without bubbles or leaks in the circuit. *SEE TROUBLESHOOTING*
3.3.10 Move the set up to the cell culture test incubator to keep the cells at the appropriate temperature. Leaves the flow running as long as necessary for experimental purposes.

**Figure 1:**
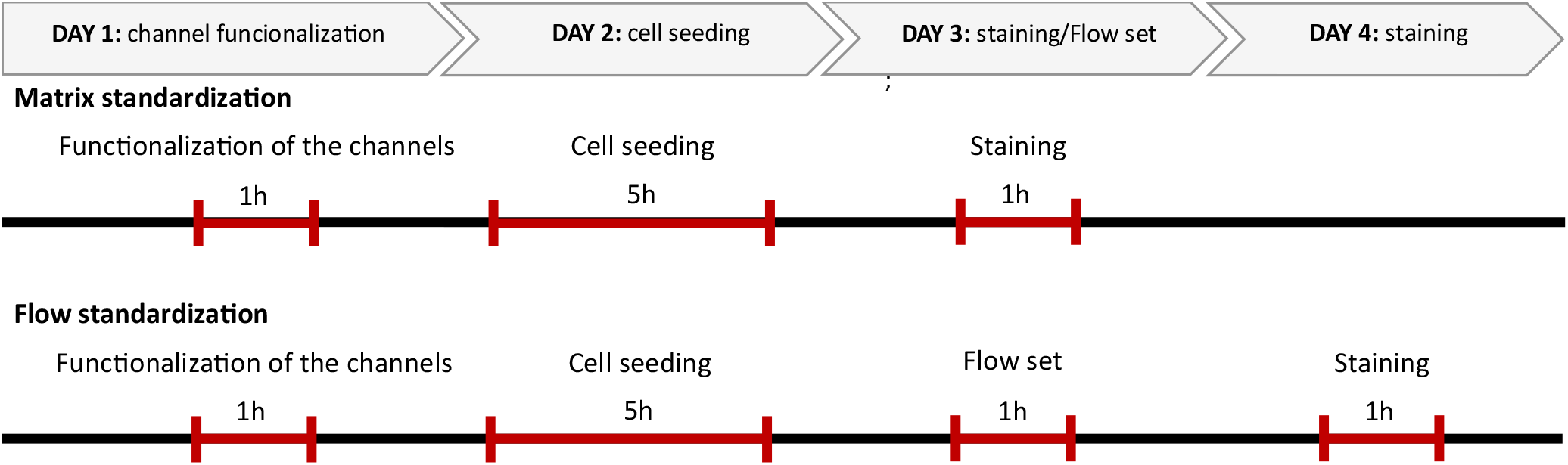
Schematic representation of the timing for both experimental settings, matrix and flow standardizations.

**Figure 2.**
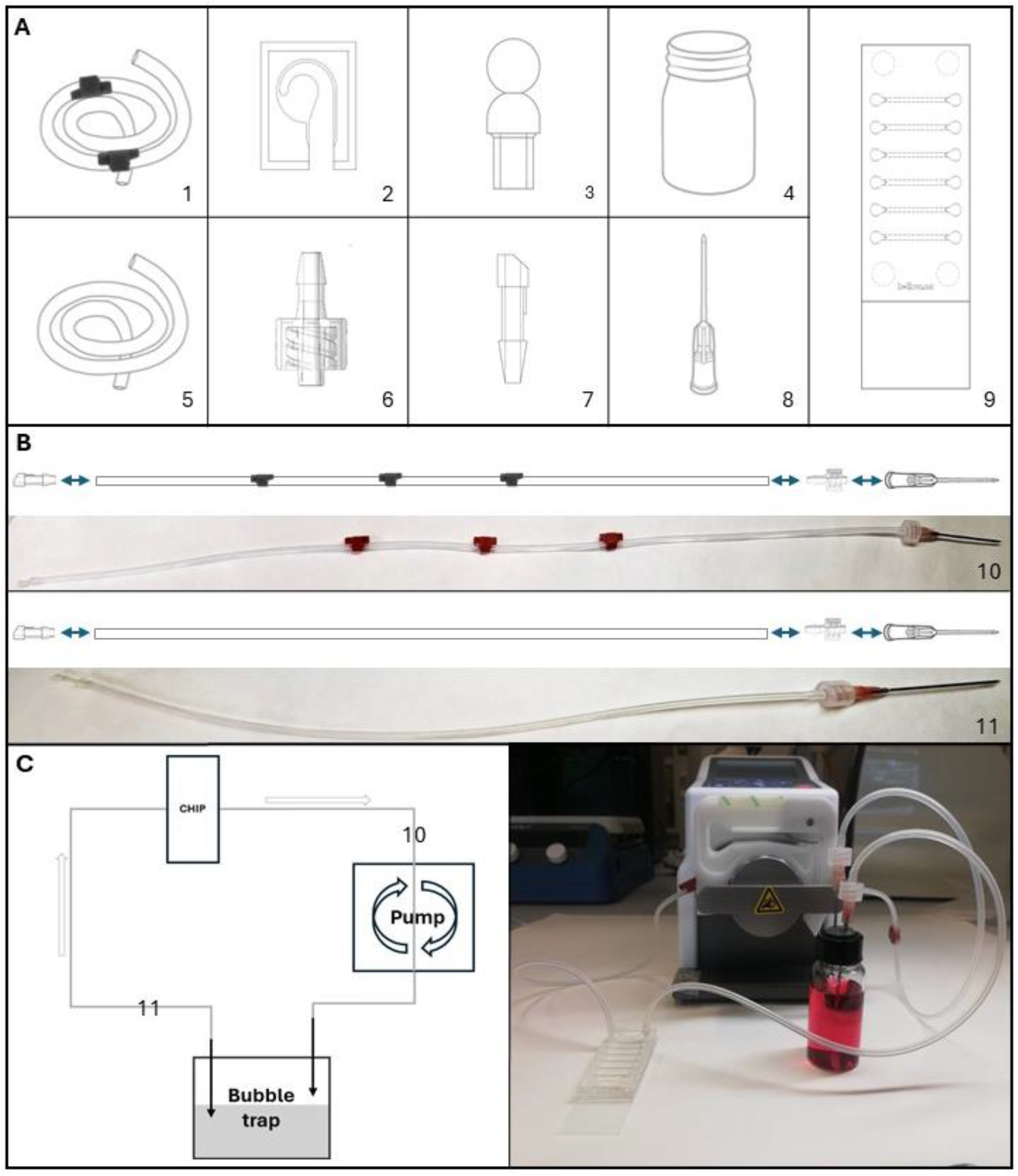
Schematic representation of the materials to set the experiment under flow. **A**) The individual materials are numbered as follows: 1-3-stop platinum-cured silicone tubing, 2-Open jaw slide clamp, 3-Straight press-in plug, 4-Reservoirs with septum cap, 5-Silicone tubing, 6-Male luer lock connector, 7-Straight chip connector, 8-Syringe needle, 9-Multi-linear channel chip. **B:** Representation of the assembly of the tubes, divided in two settings. 10-Tube from chip through the pump and to the reservoir composed by the materials 7+1+6+8 indicated in Figure 2A; 11-Tube from reservoir to the inlet composed by the materials 7+5+6+8 indicated in Figure 2A. C: Schematic representation of the system on the left (including the chip (material 9), the tube connecting the outlet of the chip and the reservoir through the pump (material 10), the bubble trap with the reservoir (material 4) and the tube from the reservoir to the inlet of the chip (material 11), the real mounted system is showed on the right.

#### BOX 2 | Bubble trap

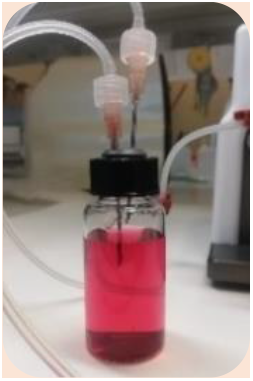

The purpose of this mechanism is to prevent bubbles from entering the microfluidic chip and disturbing the flow as well as the cells. For the bubble trap to function properly, the needle connected to the tube with stoppers coming from the pump must not be into the medium inside the reservoir, allowing the medium to drip and the needle connected to the tubing going into the chip must be submerged inside the medium to prevent air from entering into the chip.

### 3.4 LIVE CELL STAINING *TIMING 1h*

3.4.1 Remove the set up with the chip from the test incubator and take out the tubes from the channels.
3.4.2 For the cell staining, follow the steps 2.3.2 and 2.3.3.

### 3.5 IMAGE ACQUISITION

3.5.1 For image acquisition, follow the indications from 2.5.

## PROTOCOL GLOBAL TIMING

## TROUBLESHOOTING

## Anticipated results

The described protocol was applied using the indicated multi-linear channel chips (BFlow, B06_0024) with two cell lines, Caki-1 and A549, being relevant cell cultures to test toxicity in kidney and lung, respectively. For the matrix standardization, ten different matrixes were tested. Matrixes that worked in static conditions were selected to be tested under flow conditions.

### Matrix standardization of Caki-1 and A549 cell lines

A total of three biological replicates have been made for each of the matrix conditions for each cell line, analysing the number of cells using the nuclear staining in Caki-1 and A549 (Figure 3 and Figure **4**, respectively).

**Figure 3:**
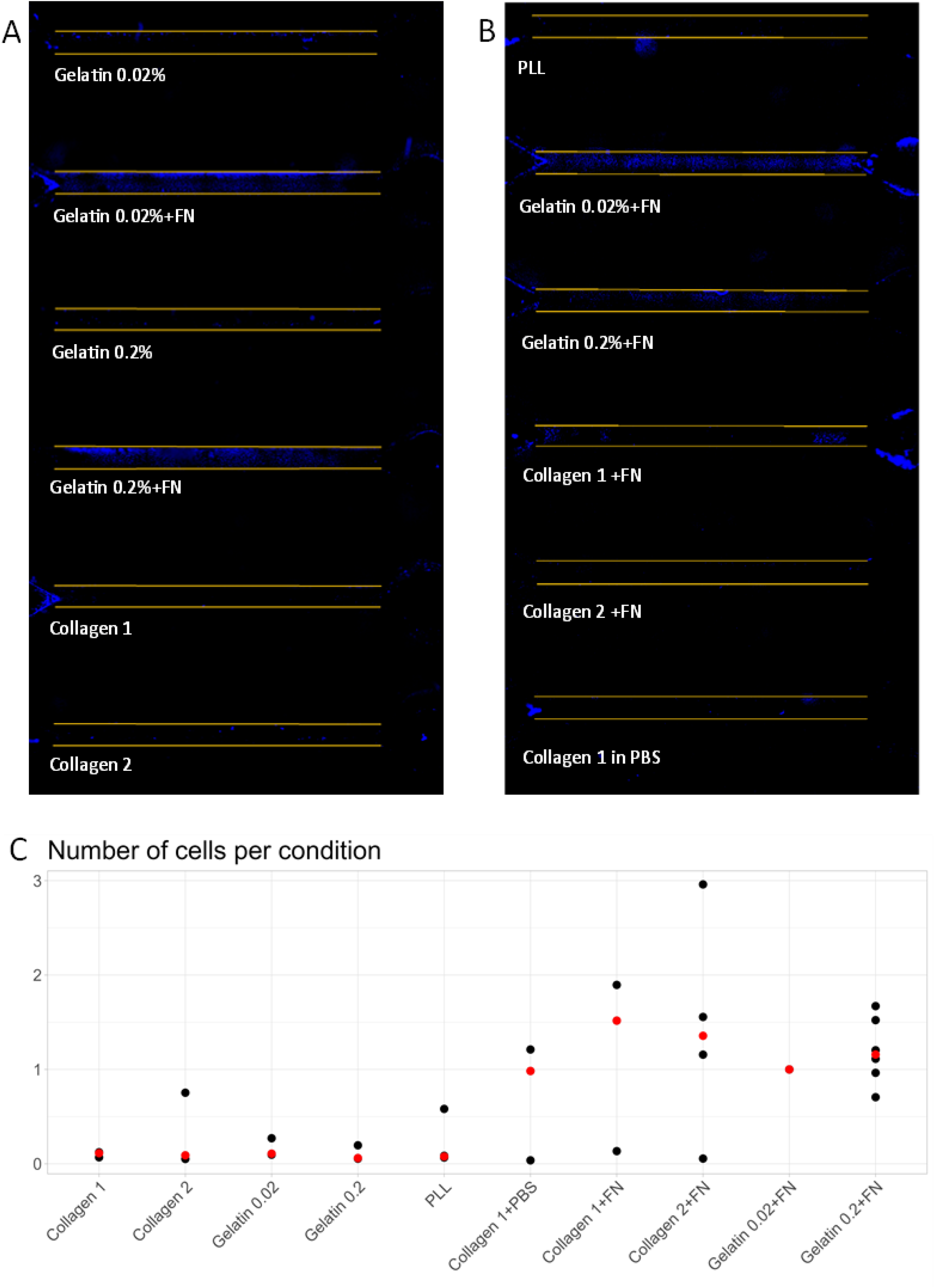
Caki-1 cell line matrix standardization. A and B) representative fluorescent microscopy of the multi-linear channel chips showing the stained cell nuclei in blue for all the tested conditions. Yellow lines are showing the edges of the channels, being 1 mm the distance between lines. C) Dot plot illustrating the effectiveness of the ten types of matrixes assessed. Relative values with gelatine 0.02% + fibronectin are showed. The red dot represents the median value.

**Figure 4:**
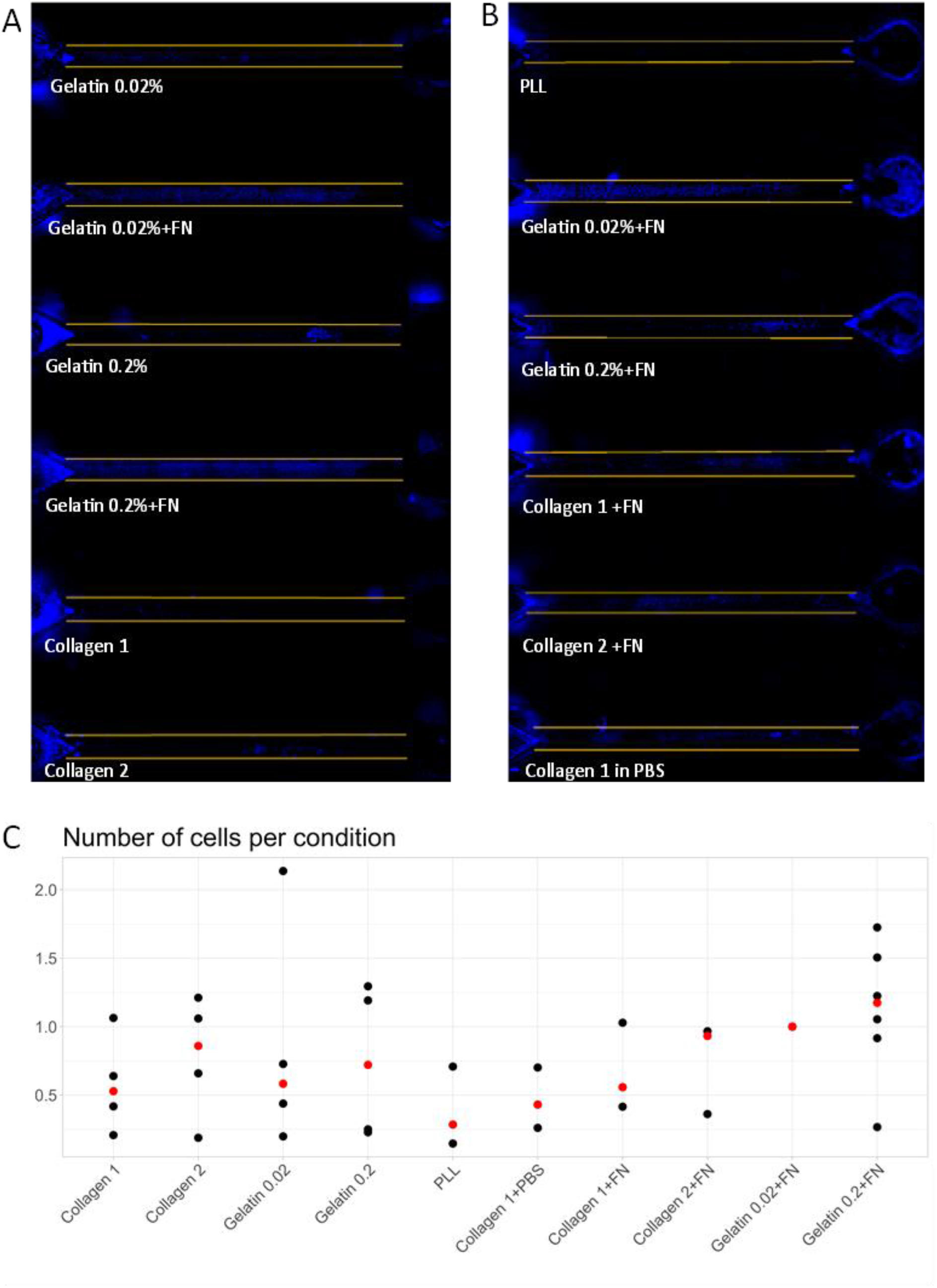
A549 cell line matrix standardization. A549 cell line. A and B) representative fluorescent microscopy images of the multi-linear channel chips showing the stained cell nuclei in blue for all the tested conditions. Yellow lines are showing the edges of the channels, being 1 mm the distance between lines. C) Dot plot illustrating the effectiveness of the ten types of matrixes assessed. Relative values with gelatine 0.02% + fibronectin are showed. The red dot represents the median value.

In Caki-1 cell culture, coating with the combination of gelatine and fibronectin (at both concentrations of gelatine) showed higher number of cells than gelatine alone (Figure 3). The combination of collagen and fibronectin (using both collagens) showed lower adhesion of cells compared with gelatine+fibronectin coating. As in the case of gelatine, the combination of collagen and fibronectin improve the adhesion compared with collagen alone. Poly-L-lysine (PLL) functionalization did not show Caki-1 cell adhesion (Figure 3). Relative values of the number of cells are represented in Figure 3C. Performing a One Sample t Test (value of 1 is gelatine 0.02%+fibronectin) showed that collagen 1 (Gibco), gelatine 0.02%, gelatine 0.2% and PLL were significantly different compared to the reference value (number of nuclei in gelatine 0.02%+fibronectin). In A549 cell culture, no clear differences were observed. Gelatine alone provided low and no reproducible adhesion (

Figure **4**). However, when fibronectin was added, 60% the experiments exhibited A549 cell adhesion (

Figure **4**). In the case of collagen coating, A549 cells showed poor adhesion with collagen alone, but upon addition of fibronectin, 50% of the experiments exhibited cell adhesion albeit with a uniform distribution. Consistent with the findings with Caki-1 cells, it appears that PLL functionalization did not exhibit cell adhesion (Figure **4**). No significant effect was observed by One Sample t Test compared with the reference value of 1 (Gelatine 0.02%+fibronectin). These observations underscore the importance of matrix selection and standardization for microenvironment improvement, particularly the inclusion of fibronectin, promoting cell adhesion for both Caki-1 and A549 cells.

### Flow standardization of Caki-1 and A549 cell lines

Based on the analysis of the previous results, it was determined that the most effective combination for channel functionalization under flow conditions is gelatine plus fibronectin for both cell types. The cell culture capability to hold flow conditions into an OoC is directly related with the quality of the cellular adhesion and a proper biological microenvironment. In this way, is important to test the critical flow levels that these cell lines can hold, stablishing the upper limit of wall shear stress for OoC experiments. To achieve effective flow standardization in organ chips, seven different flow conditions were investigated and evaluated, covering a wide range of flow rates from 160 to 5000 µl/min (table 2) which cover a range from 0.40 to 12.65 dyne/cm^2^ wall shear stress (TABLE 2).

**TABLE 1.**
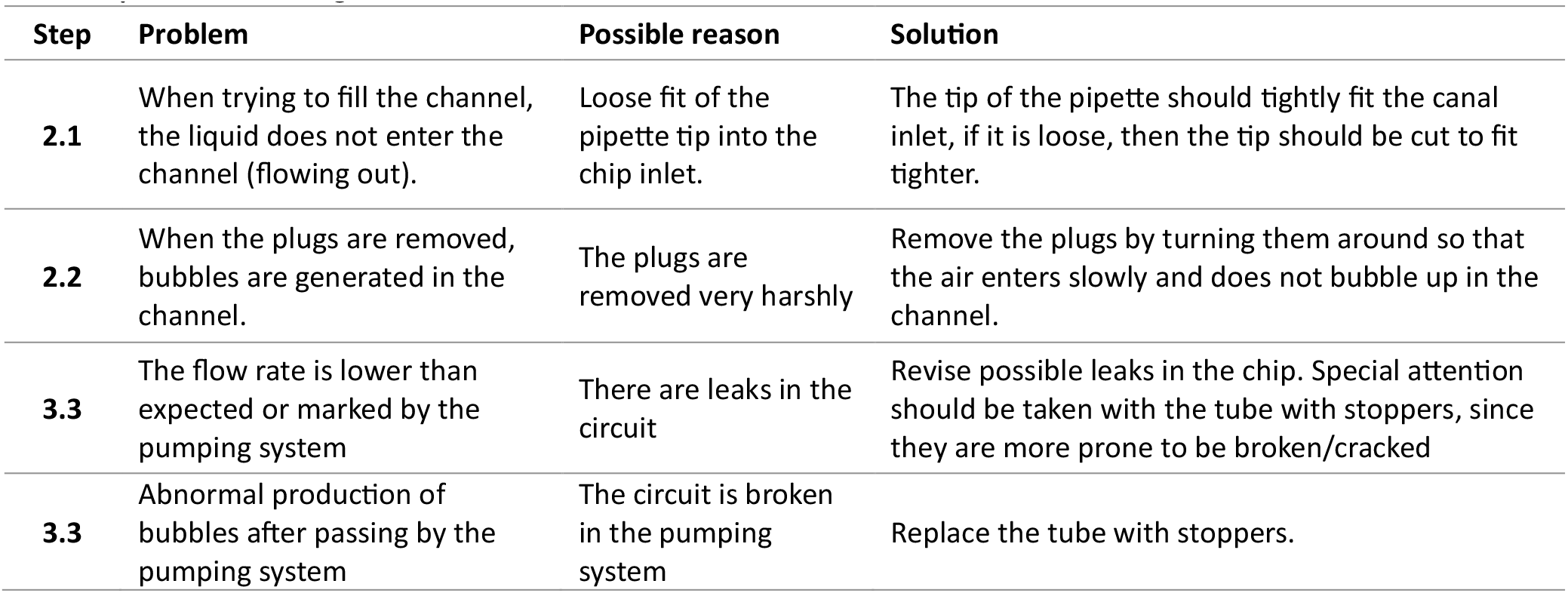
Troubleshooting table.

**TABLE 2.** Flow rate and the corresponding wall shear stress in the channels used (1 mm of diameter with circular section). The referenced images are from Figure 5 and Figure 6.

| Image | Flow rate | Wall shear stress $\tau$ |
| --- | --- | --- |
| A | 160.00 $\mu\text{L}/\text{min}$ | 0.40 $\text{dyn}/\text{cm}^2$ |
| B | 320.00 $\mu\text{L}/\text{min}$ | 0.81 $\text{dyn}/\text{cm}^2$ |
| C | 640.00 $\mu\text{L}/\text{min}$ | 1.62 $\text{dyn}/\text{cm}^2$ |
| D | 1.20e+3 $\mu\text{L}/\text{min}$ | 3.04 $\text{dyn}/\text{cm}^2$ |
| E | 2500 $\mu\text{L}/\text{min}$ | 6.32 $\text{dyn}/\text{cm}^2$ |
| F | 5000 $\mu\text{L}/\text{min}$ | 12.65 $\text{dyn}/\text{cm}^2$ |
| G | 0 $\mu\text{L}/\text{min}$ | 0 $\text{dyn}/\text{cm}^2$ |
| H | STATIC | - |

A comprehensive evaluation of the flow conditions yielded significant insights into the behaviour of both cell lines in response to these flow variations. Across all tested conditions, it was observed that both the Caki-1 (Figure 5) and the A549 (Figure 6) line remained adhered to the channels walls over all the flow tested, suggesting robustness of this microfluidic system in maintaining cellular integrity under different flow conditions. Of particular interest was the phenomenon observed regarding the nuclear morphology of Caki-1 cells. A notable trend towards rounding of the cellular nucleus in the presence of flow was noted (data not shown), suggesting an adaptive response of cells to the shear forces generated by flow.

**Figure 5:**
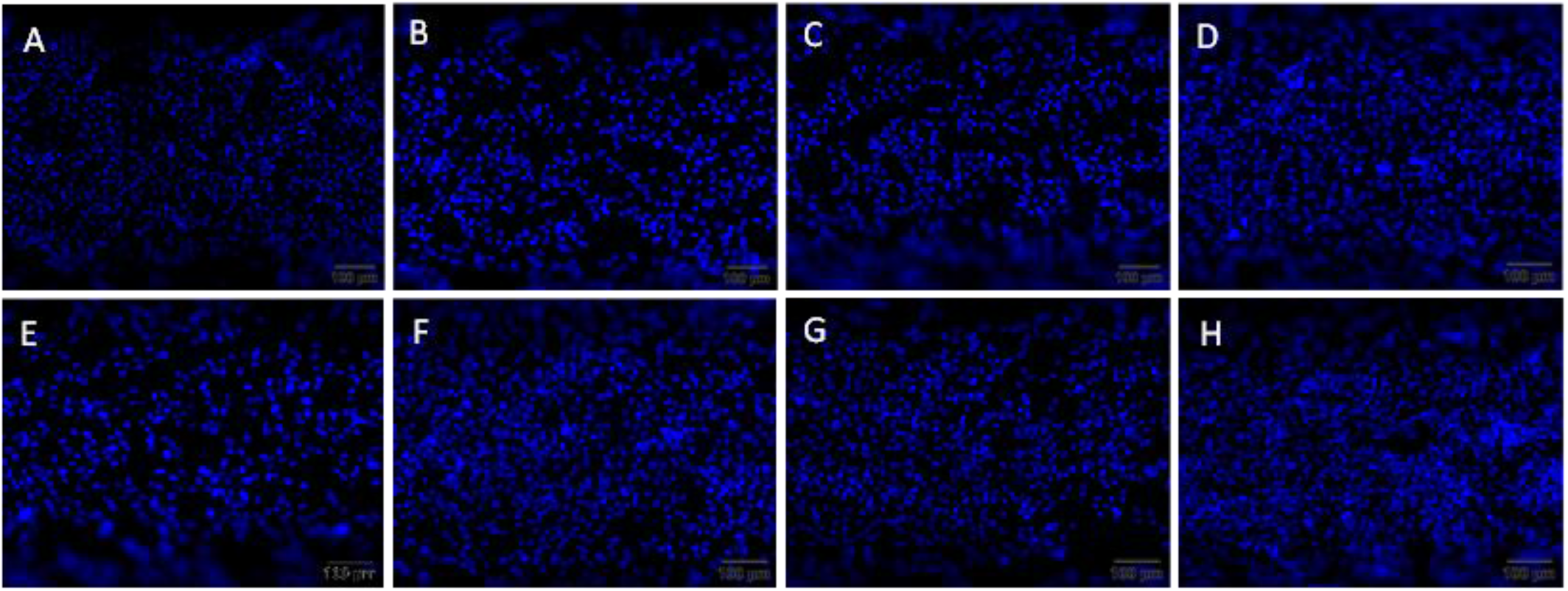
Caki-1 cell line under flow. Representative fluorescent microscopy images of the multi-linear channel chips showing the stained cell nuclei in blue for all the tested conditions in µL/min: A) 160, B) 320, C) 640, D) 1200, E) 2500, F) 5000, G) 0, H) static (as indicated in Table 2). The scale bar is 100 µm.

**Figure 6:**
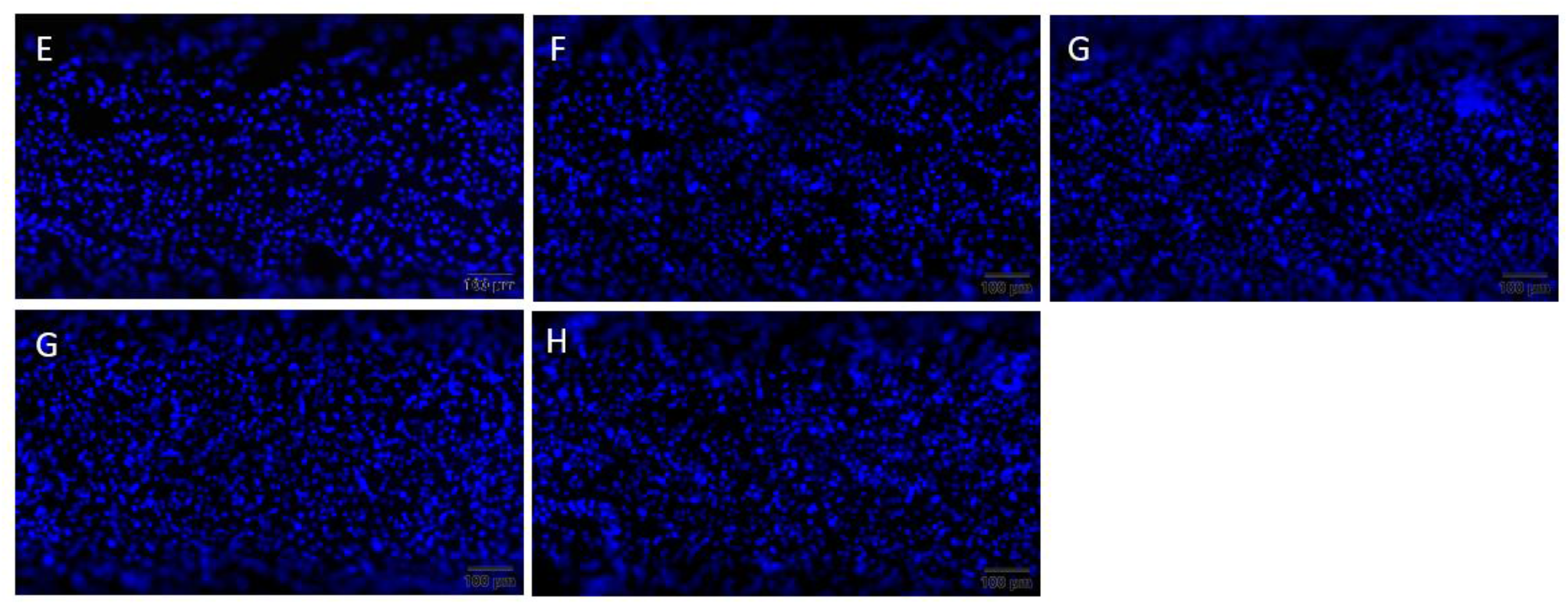
a549 cell under flow. Representative fluorescent microscopy images of the multi-linear channel chips showing the stained cell nuclei in blue for all the tested conditions in µL/min: E) 1200, F) 2500, G) 5000, H) 0, I) static (as indicated in table 2). The scale bar is 100 µm.

These findings not only highlight the ability of both cell types to hold a broad set of flow conditions, but also provide valuable insights into how the microfluidic environment may modulate cellular morphology and response. The protocol indicated in this work aims to guide the OoC user in the rational standardization to understand the limits and conditions of new OoC systems. This systematic approach provides insight into how different flow rates can influence cellular responses and system dynamics, being applicable in other combinations of microfluidic channels, cellular cultures and extracellular matrix coatings.

## Final considerations

In summary, three main considerations must be addressed for the implementation of experimental procedures in OoC systems under flow conditions.

First, the selection of the ECM or surface coating of the chip must be tailored and optimized for each specific cell type employed. It is evident that each cell type exhibits varying adhesion capacities, and not all materials or surfaces are suitable for cell culture. Given that cells in these systems will be subjected to wall shear stress from fluid flow, it is crucial to ensure robust cell adhesion and maintenance throughout the experiments. As demonstrated in this study, incorporating fibronectin into the surface coating can serve as a general guideline for enhancing cell adhesion across various cell types.

Second, it is essential to establish the appropriate biological microenvironment following cell adhesion to the chip. This can be achieved by optimizing several parameters, including allowing adequate time for cell differentiation/maturation into the chip, using a suitable culture medium, and applying a low flow of medium before exposing the cells to the final experimental conditions. These steps are vital to ensure optimal cell performance during subsequent experimental procedures.

Finally, the objective of OoC experiments is to analyse the behaviour and response of the cells within a novel microenvironment under flow conditions. This needs the measurement and consideration of new parameters to evaluate cell functionality accurately. Additionally, it is important to consider that these experiments may yield new insights into the responses of cells previously studied under static conditions.

## Acknowledgements

Andrea Balsa-Díaz, Laura Vázquez-Vázquez and Lois Rivas-Meizoso contracts were funded by Galician Innovation Agency (GAIN) INVESTIGO program. Elías Ferreiro-Vila and Bruno Kotska Rodiño-Janeiro contracts were funded by National Research Agency Torres Quevedo program 2022 and 2020 (PTQ2022-012762 and PTQ2020-011081, respectively). The work was partially funded by Center for Technological Development and Innovation NEOTEC 2022 program (SNEO-20222232), School of Industrial Organization ActivaStartUP program (ATS_PYME_004#36), “La Caixa” Foundation Caixa Impulse Validate 2021 Call (CI20-00200), Xunta de Galicia (ED431B 2023/07), GAIN CONECTA COVID program (IN852E 2021/2), and the grant PID2022-138322OB-100 funded by MCIN/AEI/10.13039/501100011033.

